# Activation of Transposable Elements Upon Statin Treatment

**DOI:** 10.1101/2022.05.01.490198

**Authors:** Franco Valdebenito-Maturana, Braulio Valdebenito-Maturana

**Affiliations:** Facultad de Farmacia, Barrio Universitario s/n, Concepción, 4030000, Chile; Núcleo Científico Multidisciplinario, School of Medicine, Universidad de Talca, Talca, Chile

**Keywords:** Transposable Elements, Gene regulation, Statins, Pharmacogenomics

## Abstract

High cholesterol levels have been associated with cardiovascular diseases, and lowering them has been a key focus in the treatment of such diseases. Statins are drugs used with that aim, and can be divided in the lipophilic Simvastatin and the hydrophilic Rosuvastatin. Regardless of the statin type, a high proportion (∼70%) of patients stop using statins due to suffering from side effects on skeletal muscle, such as myalgia, and muscle cramps. Thus, there has been a considerable effort in understanding how statins contribute to these side effects. A catalogue of genes and molecular pathways that change upon statin treatment has been recently published, allowing further understanding how the side effects occur. However, Transposable Elements (TEs) were not studied. TEs can move within a genome, and they are highly repetitive, representing about half of the human genome. Currently, most TEs in the human genome are inactive, but it has been shown that TEs can still transcribe, and that either via their transposition or their transcriptional activity, can influence gene expression. Here, using novel computational tools to accurately estimate TE expression, we studied their activity and predicted their potential impact on gene expression. We developed a catalogue of TEs expressed upon statin treatment, and the putative genes whose expression might be influenced by TEs. Overall, we speculate that based on our findings, TEs might be a key target in order to understand statin-mediated side effects.

## 1. Introduction

Cardiovascular diseases are one of the most common causes of death in developed countries ^1^. High low-density lipoprotein (LDL) cholesterol levels are one of the key culprits in the development of these pathologies ^2^, and lowering its levels has been a main focus of treatments ^3^. Nonetheless, adherence to medical treatments is often conditioned to the extent of the patient’s tolerance to side effects ^4^. One of the most used drugs to decrease LDL cholesterol levels are statins ^5^. There are several types of statins, with the following two types of statins being commonly studied: Simvastatin and Rosuvastatin, whose main difference lies in that the former is lipophilic, whereas the latter is hydrophilic. Regardless of the statin type, it has been estimated that around 70% of the patients using these drugs, show pervasive side effects on skeletal muscle, and thus stop taking them^6^.

Statin-induced myopathy is often presented in the form of muscle cramps, pain (myalgia) and weakness. A catalogue of differentially regulated genes in human cells, upon statin treatments revealed extensive changes associated with key cellular process, such as RNA metabolism and pain signaling pathways ^6^. However, the expression of Transposable Elements was not analyzed.

Transposable Elements (TEs) have the genetic machinery to move within a genome, and thus, are part of the repetitive portion of many genomes, representing almost half of the human and mouse genomes. Depending on their mechanism of transposition, they can be divided into the DNA class or the Retrotransposon class. DNA TEs produce proteins that “cut” the element from its original genome locus and insert it elsewhere, whereas Retrotransposons perform reverse transcription on their RNAs, resulting in a new copy of the element, which is later inserted in another location. Moreover, Retrotransposons are further classified into Long Terminal Repeats (LTRs), the Long Interspersed Nuclear Elements (LINEs) and Short Interspersed Nuclear Elements (SINEs) ^7^. Regardless of their class and type, most TEs are not able to transpose due to either accumulation of mutations, or to epigenetic silencing. Current evidence shows that despite this, TEs can transcribe and this in turn might play key roles in gene regulation. Thus, TEs either by the transcriptional or transpositional activity might impact gene expression ^8^. Examples of processes in which TEs have been linked to regulation are mammalian development, and in disease ^9,10^. As TEs are repeated in a genome, they are often discarded in regular gene expression studies, impeding the study of their role. Specifically, their role in the genetic response associated to side effects to drug treatments has not been studied. Perhaps the most interesting example of this is a recent work showing that TEs were involved in the response to insecticides in the fruit fly ^11^. Thus, this would suggest that TEs might play a role in the cellular stress that occurs as response to pharmacological interventions.

The aim of this work was to (1) identify if TEs become differentially regulated as response to Simvastatin and Rosuvastatin treatment, and (2) via stringent statistical analysis associate TEs with potential mechanisms of gene regulation associated to the side effects of statins.

## 2. Methods

RNA-Seq data corresponding to untreated, Simvastatin treated and Rosuvastatin treated samples were previously published by Grunwald et al. (2020). The datasets were downloaded from the public SRA database, accession SRP126593. Mapping and quantification of reads per gene and TEs was done using SQuIRE ^12^, which employs technical advances that allow the estimation of TE expression in a locus-specific manner. Differential expression analysis where done with DESeq2 ^13^, using the Simvastatin or Rosuvastatin treated samples versus the untreated samples. A threshold of |log_2(FC)| ≥1.5 and adjusted P-value ≤ 0.05 were used as criteria for significant differential expression.

Locus-specific analysis of TEs was done in several steps using BEDTools ^14^. First, TEs overlapping with genes were classified into “Exon”, “Start codon”, “Stop codon”, or “Intronic” depending on which of these elements they had an overlap. TEs fully covered by exons, start or stop codons were discarded, as these cases probably correspond to the entire gene being expressed rather than the TE. TEs not overlapping any gene were classified as “Intergenic”. Intergenic TEs were associated with their closest downstream gene.

Statistical association between TEs and genes was done using TEffectR ^15^. TEffectR applies a linear modeling in which the response variable is the gene expression, and the independent variable is the TE expression. This allows the statistical prediction of whether changes in gene expression can be explained by changes in TE expression. Gene-TE associations with p-value ≤ 0.05 were selected. Afterwards, the correlation between the Gene-TE pairs was assessed with the R environment for statistical computing ^16^. To explore the impact of TE expression in gene regulation, pairs with negative correlation were selected. STRING database v11.0b ^17^ was used for gene network analysis and enrichment, using statistical threshold of FDR ≤ 0.05.

All plots were done using ggplot2 ^18^.

## 3. Results

### 3.1. Transposable Elements become differentially expressed as a response to Statin treatment

Human cells treated with Statins show differential expression of several genes, which may explain the molecular underpinnings associated to the side-effects provoked by their use. Transposable Elements were not studied previously due to technical difficulties associated to their expression quantification from RNA-Seq data ^6^. To address this, quantification of TE expression from was performed using SQuIRE. This tool allows the locus-specific analysis of TEs, which is the first step towards understanding and predicting the potential impact of TEs in their genomic vicinity. After the utilization of SQuIRE, differential expression analysis of TEs was performed, revealing that a large number of TEs show changes in their expression upon Statin treatment (Fig 1).

**Fig 1.**
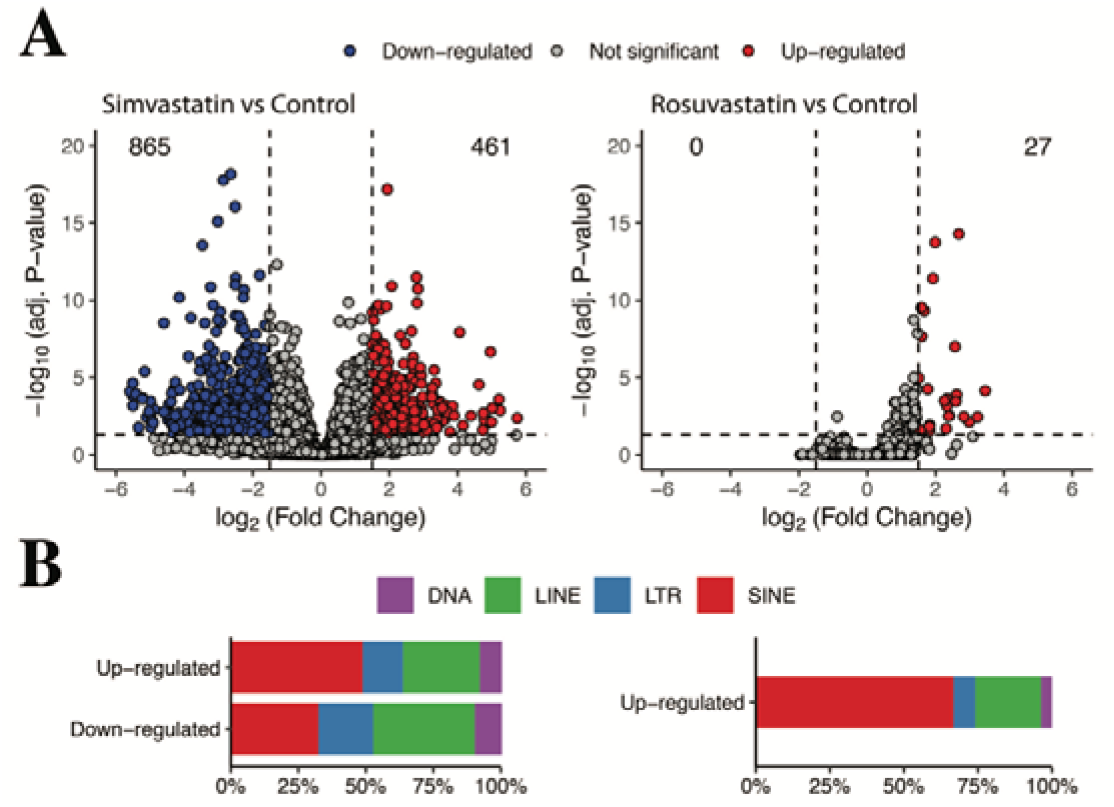
A. Volcano plots of Transposable Element differential expression. Left, comparison of Simvastatin vs Control samples, Right, comparison of Rosuvastatin vs Control samples. In each plot, red points correspond to up-regulated TEs (increased expression in Simvastatin or Rosuvastatin relative to Control, respectively), blue points correspond to down-regulated TEs, and grey points correspond to TEs with no statistically significant differences. Horizontal dashed line corresponds to adjusted P-value 0.05, and vertical dashed lines correspond to the -1.5 and 1.5 fold change threshold. Numbers at the upper left of each plot correspond to down-regulated TEs, and those at the upper right to up-regulated TEs. B. Class distribution of up-regulated and down-regulated TEs in Simvastatin (left) or in Rosuvastatin (right).

Depending on which statin was used, the number of differentially expressed TEs is significantly different. A total of 1326 TEs changed their expression upon Simvastatin treatment, whereas only 27 did upon Rosuvastatin treatment. Moreover, in the latter, the 27 TEs became up-regulated, while in the former 461 (34.8%) were up-regulated and 865 (65.2%) down-regulated (Fig 1A). Interestingly, in all cases there is a predominance of SINE TEs, followed by LINE TEs (Fig 1B). It is worth noting that SINE TEs are known to negatively influence several mRNA associated processes such as export from the cytoplasm, and translation ^19^.

### 3.2. Transposable Elements are significantly associated to genes

After finding differentially expressed TEs, our next aim was to predict their potential role in gene regulation. To do so, we employed a rigorous statistical approach. First, we used TEffectR to identify the genes whose expression could be explained by changes in TE expression. Afterwards, we only kept the gene-TE pairs having statistical significance according to TEffectR (p≤0.05). For these significant gene-TE pairs, we then calculated their pairwise correlations. The pairwise correlations were then plot according to the Statin used (Fig 2).

**Fig 2.**
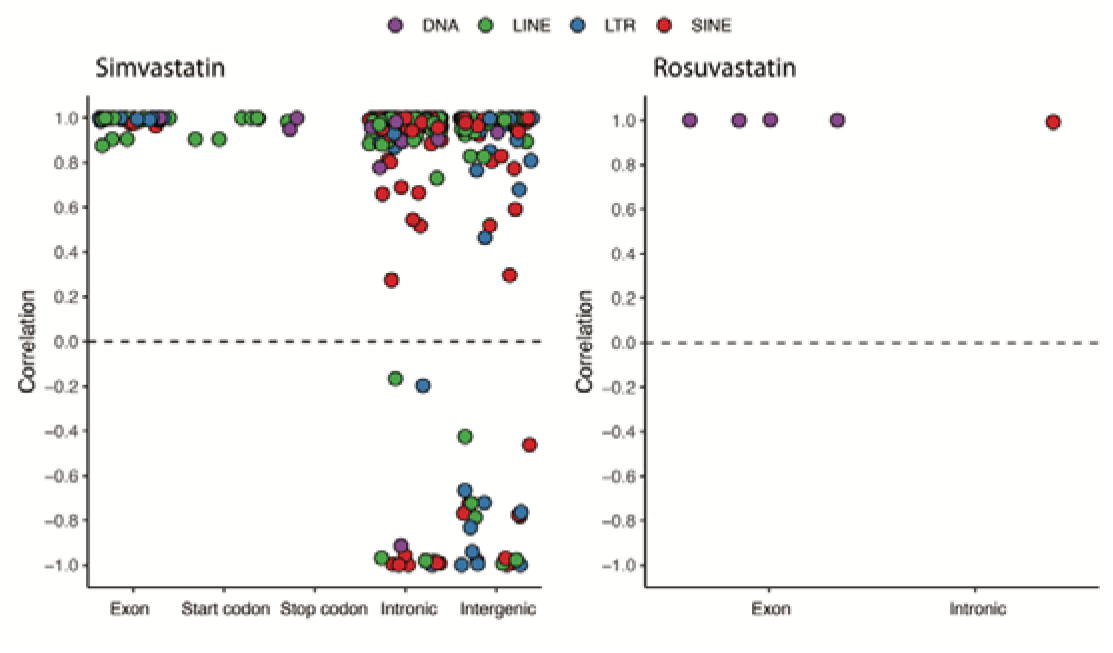
Gene-TE correlation distribution plots according TE classes. Left: correlations amongst Simvastatin gene-TE pairs, Right: correlation distribution amongst Rosuvastatin gene-TE pairs. Dashed line at 0 was added to aid in the visual separation of positive and negative correlations.

Most of the correlations were positive, suggesting that most expressed TEs are probably expressed as a consequence of its host gene being expressed. This phenomenon, known as TE co-transcription is not well understood, and it is not really clear how it might impact gene regulation. On the other hand, Intergenic TEs with positive correlation to their closest downstream gene might indicate that these TEs could be acting as alternative transcription start sites, or as enhancers, depending on how far they are to their respective genes. Interestingly, there are several Intronic and Intergenic TEs that correlate negatively with their associated gene. For the next section, we only focused on these TEs with negative correlations, as we argue that these are the most likely to play a bigger role in Statins side-effects.

### 3.3. Transposable Elements might influence the expression of key genes upon Statin treatment

After our stringent statistical analysis, we identified 15 genes negatively correlated with intronic TEs (Table 1), and 21 negatively correlated with intergenic TEs (Table 2).

**Table 1.**
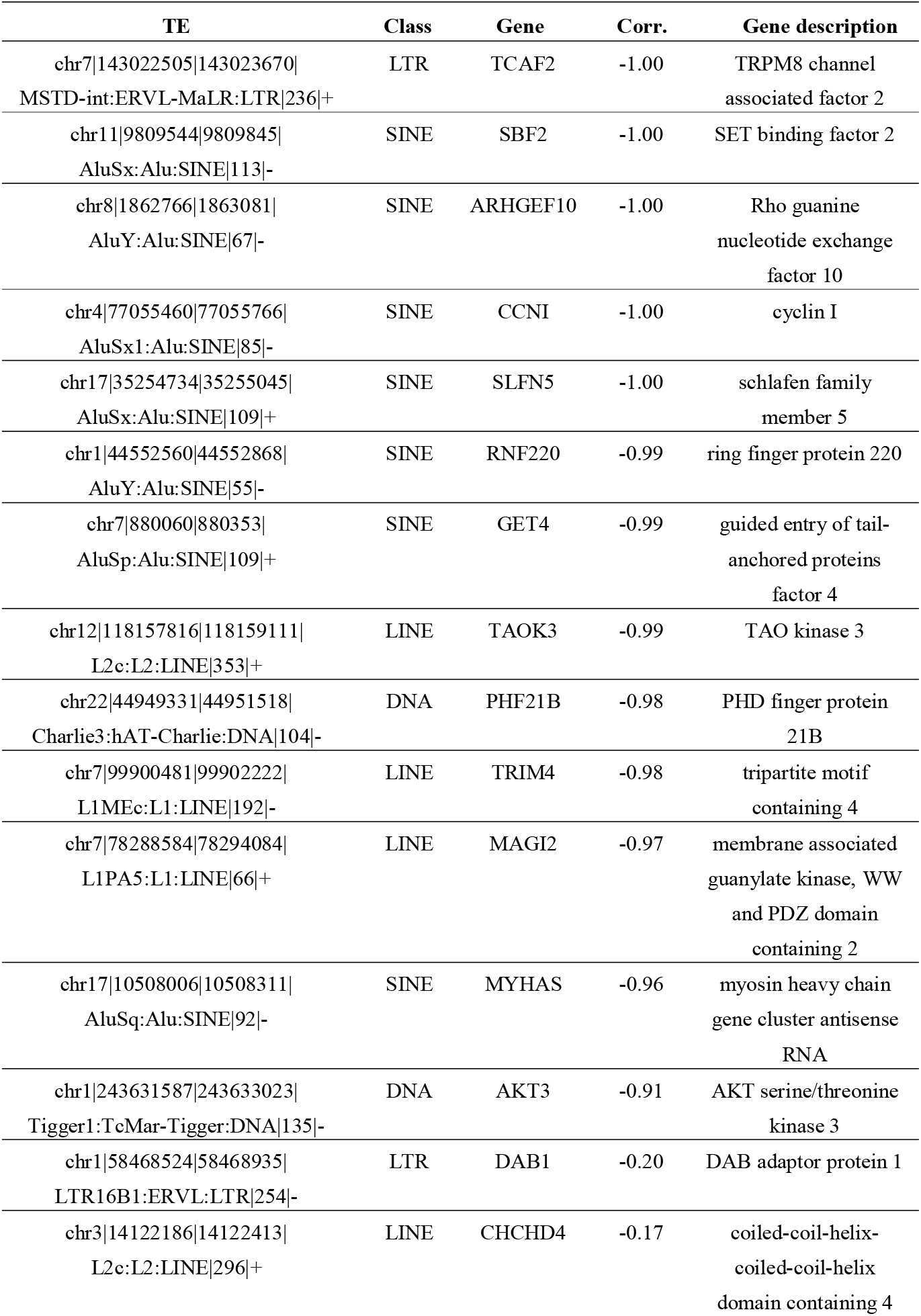
Intronic TEs negatively correlated with their associated genes.

**Table 2.**
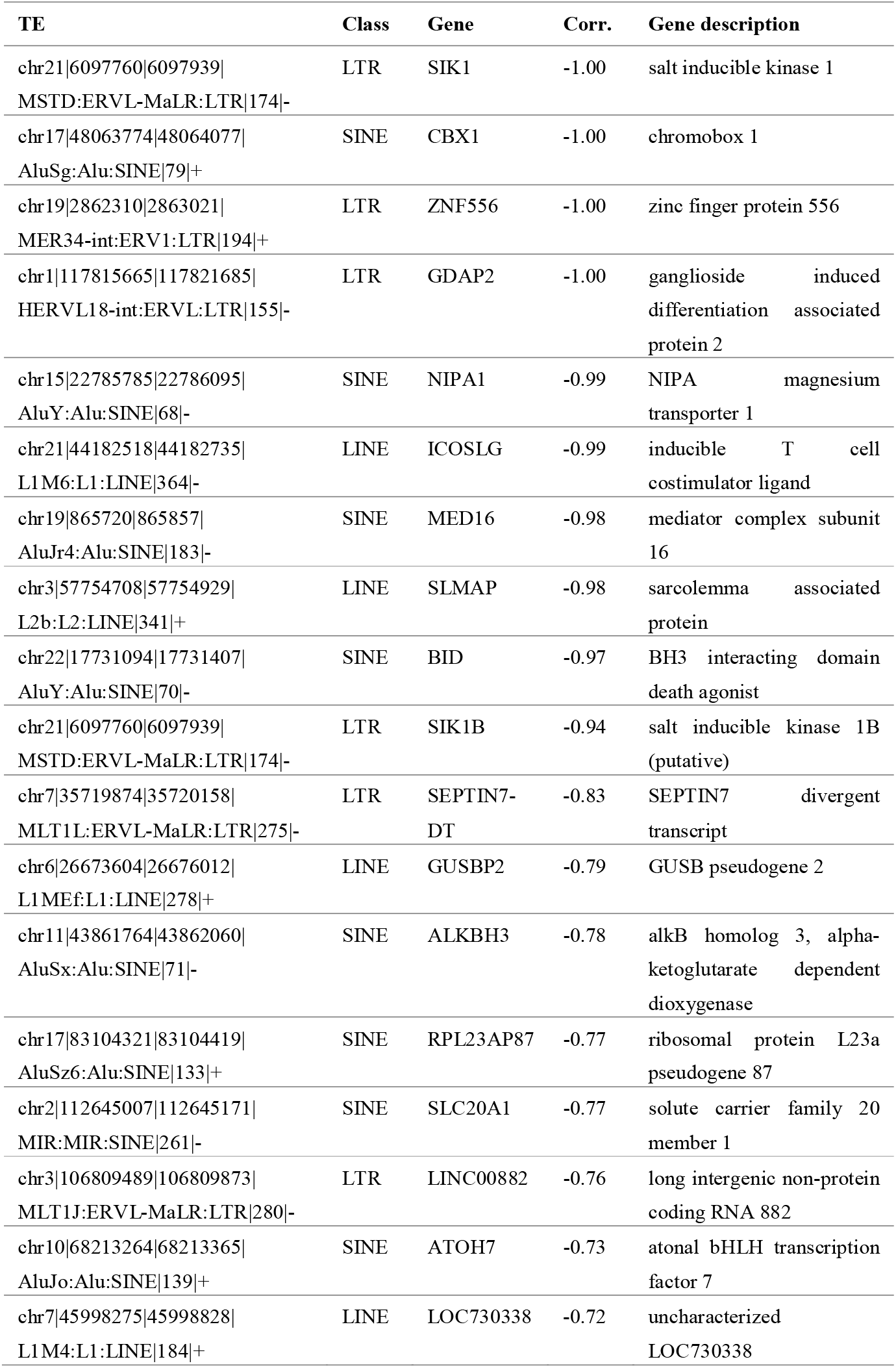
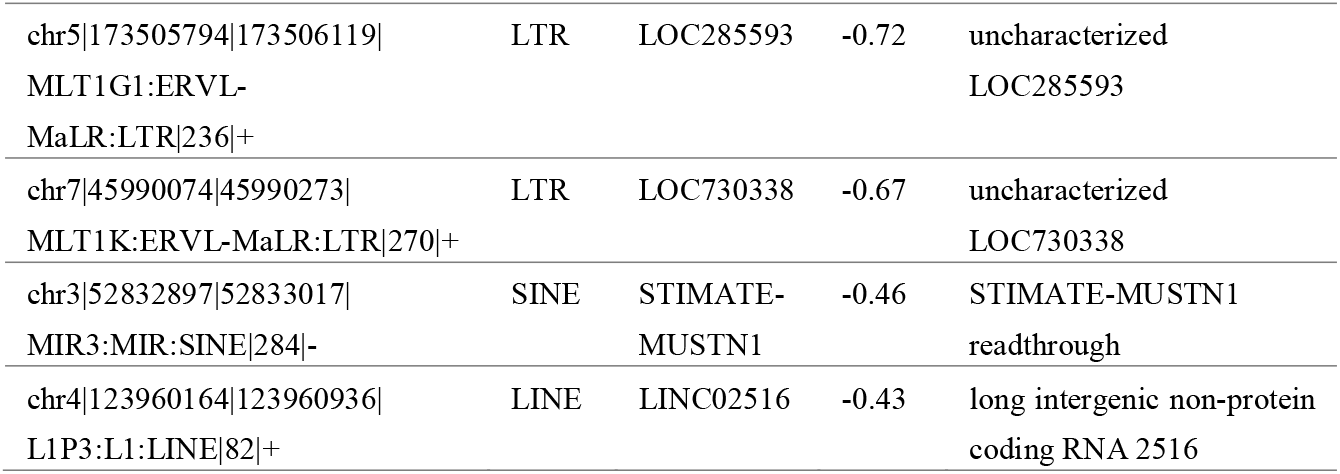
Intergenic TEs negatively correlated with their associated genes.

Amongst all the genes having negative correlations with TEs, we first highlight ZNF556 (zinc finger protein 556), ATOH7 (atonal bHLH transcription factor 7), TAOK3 (TAO kinase 3), AKT3 (AKT serine/threonine kinase 3). Zinc finger proteins are known to play important roles in cellular processes, such as transcriptional regulation ^20^. At the same time, transcription factors are master regulators of other genes ^21^. Moreover, TAO kinase 3 has been linked to the MAPK14 stress activated molecular pathway, and AKT serine/threonine kinase 3 is involved in cell signaling and glucose uptake ^22^. Collectively, this result suggests that TEs might influence the regulation of several genes that are themselves the controllers of several further genes, in turn indicating that TEs might play a role in modulating gene expression at a much larger scale. We also pay special attention to AKT3, since it has been reported that in some patients treated with Statins, defects in glucose metabolism occur ^23^.

Perhaps the most relevant gene putatively regulated by a TE, is SLMAP (sarcolemma associated protein). SLMAP might have a role in myoblast fusion ^17^, a key cellular process for adequate skeletal muscle normal development and repair ^24^. To further explore the protein interacting network of SLMAP, we used the STRING database (methods) (Fig 3). The network is shown in the standard output of STRING (Fig 3A), and with SLMAP associated proteins belonging to the TGF-Beta pathway highlighted in red (Fig 3B). We chose to highlight the TGF-Beta associated proteins due to the link between that pathway and myopathies (further discussed in the next paragraph).

**Fig 3.**
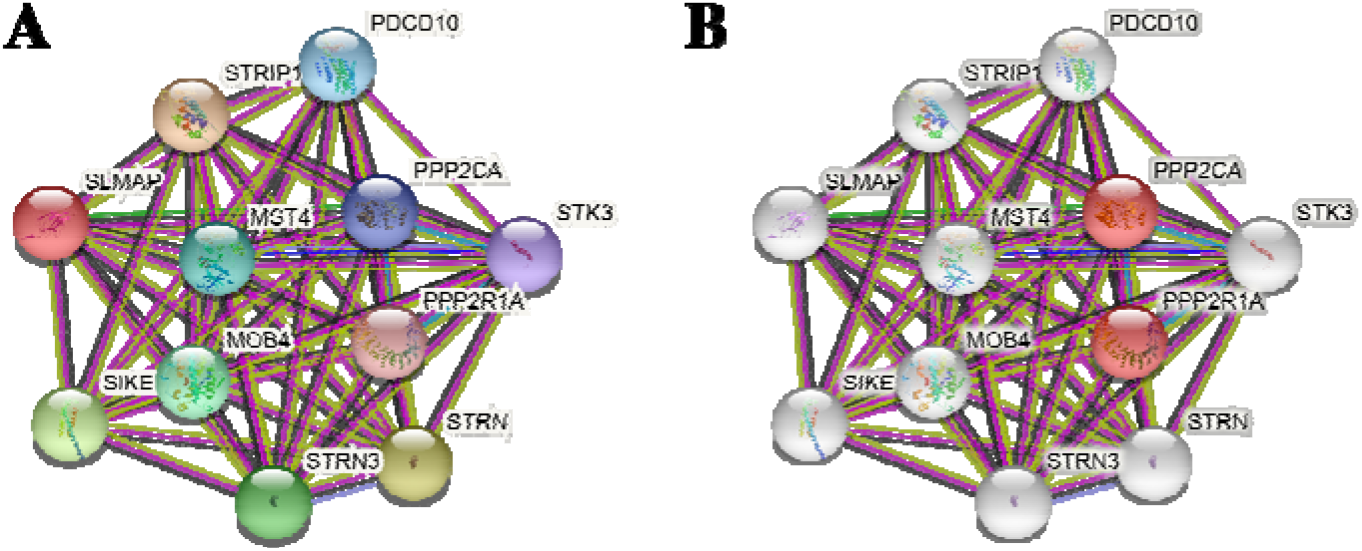
STRING network analysis. All proteins associated with SLMAP correspond to direct associations (i.e., immediate neighbors, the first shell of interaction reported by STRING) A, full network; B, full network with nodes highlighted in red according to their role in the TGF-Beta pathway. Lines connecting the nodes are colored in 3 categories: (1) known interactions: light blue from curated databases and purple for experimentally determined interactions; (2) predicted interactions: green, gene neighborhood and blue for gene co-occurrence; and (3) others: light green, text mining and black, co-expression.

We found PPP2CA and PPP2R1A as direct neighbors of SLMAP, and they both are part of the TGF-beta signaling pathway (Fig 3B). Albeit small (only 2 direct neighbors of SLMAP), according to STRING analysis, this enrichment was found to be statistically significant (FDR = 0.0063). Activation of this pathway has been implicated in skeletal muscle myopathies ^25^. Particularly, its role in myopathies is through its up-regulation. Thus, the role in TE expression related to SLMAP, and in turn, statin side-effects on skeletal muscle seem to be two-fold: (1) by impairing SLMAP normal activity, and (2) by affecting the regulatory network in which SLMAP is involved. Thus, if TE expression is effectively hindering SLMAP normal activity and its downstream targets, this could in turn be the most relevant example of TE-mediated gene regulation associated to side-effects upon Statin treatment.

## 4. Conclusions

Here we reported the first catalogue of TEs expressed upon Statin treatment. Interestingly, there are differences in terms of amounts of TEs differentially expressed depending on the Statin used, and as a consequence of this, in the amounts of TEs that can be statistically associated to genes. As reported previously, Rosuvastatin seems to have less side effects than Simvastatin. Thus, if TEs are responsible for these side effects, or mediate them, our work adds another layer of evidence in elucidating the genetic cascade that occurs upon Statin treatment. We argue that our findings will help the community interested in studying Statin side effects, and propose that TEs can be a key target in ameliorating them.

